# Model testing for distinctive functional connectivity gradients with resting-state fMRI data

**DOI:** 10.1101/298752

**Authors:** Jonathan F. O’Rawe, Jaime S. Ide, Hoi-Chung Leung

**Affiliations:** Integrative Neuroscience Program, Department of Psychology, Stony Brook University, Stony Brook, NY, 11794-2500, USA; Department of Psychiatry, Yale University School of Medicine, New Haven, CT, 06519, USA

**Keywords:** functional gradient, resting-state fMRI, striatal organization

## Abstract

In accordance with the concept of topographic organization of neuroanatomical structures, there is an increased interest in estimating and delineating continuous changes in the functional connectivity patterns across neighboring voxels within a region of interest using resting-state fMRI data. Fundamental to this functional connectivity gradient analysis is the assumption that the functional organization is stable and uniform across the region of interest. To evaluate this assumption, we developed a model testing procedure to arbitrate between overlapping, shifted, or different topographic connectivity gradients across subdivisions of a structure. We tested the procedure using the striatum, a subcortical structure consisting of the caudate nucleus and putamen, in which an extensive literature, primarily from rodents and non-human primates, suggest to have a shared topographic organization of a single diagonal gradient. We found, across multiple resting state fMRI data samples of different spatial resolutions in humans, and one macaque resting state fMRI data sample, that the models with different functional connectivity gradients across the caudate and putamen was the preferred model. The model selection procedure was validated in control conditions of checkerboard subdivisions, demonstrating the expected overlapping gradient. More specifically, while we replicated the diagonal organization of the functional connectivity gradients in both the caudate and putamen, our analysis also revealed a medial-lateral organization within the caudate. Not surprisingly, performing the same analysis assuming a unitary gradient obfuscates the medial-lateral organization of the caudate, producing only a diagonal gradient. These findings demonstrate the importance of testing basic assumptions and evaluating interpretations across species. The significance of differential topographic gradients across the putamen and caudate and the medial-lateral gradient of the caudate in humans should be tested in future studies.

## Introduction

A fundamental goal in cognitive neuroscience is characterizing the functional organization of a region (e.g. prefrontal cortex, striatum) (Kaas, 1997). In neuroimaging, global functional organization is often examined with the conventional parcellation-correlation approach, where signal from a single determined parcel (the “seed” region) is averaged across the voxels within that parcel and correlated to voxels in the rest of the brain (Hacker, Perlmutter, Criswell, Ances, & Snyder, 2012). To extend this approach to study a particular structure’s functional organization, previous studies have used a number of small seeds distributed across the structure or clustered voxels into roughly more independent subregions (e.g. Manza, Zhang, Li, & Leung, 2016; Choi, Yeo, & Buckner, 2012). Despite the theoretical significance (Jbabdi, Sotiropoulos, & Behrens, 2013; Patel, Kaplan, & Snyder, 2014; Thivierge & Marcus, 2007) investigators have only recently started to estimate spatially continuous changes in functional connectivity patterns across neighboring voxels within a parcel (Haak, Marquand, & Beckmann, 2017; Margulies et al., 2016). We refer to this approach as functional gradient mapping. In particular, Haak et al.’s (2017) method attempts to infer the functional gradient maps of a parcel (e.g. motor cortex) by computing the similarity matrix between connectivity fingerprints of all voxels in the parcel and performing spatial statistics across the manifold of the similarity matrix. One fundamental assumption made in this method is that there are no sharp shifts in topographic gradients within the parcel under investigation, implying that the parcel only has spatially contiguous functional gradients. This assumption is potentially problematic, especially for relatively confined structures that have known subdivisions of different anatomical connections (e.g. thalamus, striatum).

Here we use the striatum as the parcel of interest to describe a set of testing procedures used for examining the unitary-gradient assumption in previous models. The organization of the striatum, based on rigorous tract tracing studies in the nonhuman primate brain (Haber, 2003; Haber, Fudge, & McFarland, 2000) is suggested to follow a diagonal gradient organization from dorsolateral to ventromedial, across the entire posterior-to-anterior striatum; this striatal gradient includes both the caudate nucleus and putamen, with the internal capsule considered a relatively arbitrary division (given cortical terminal fields vary smoothly across the internal capsule). This organization was determined by examining cortical projections, with ventromedial portions of the striatum receives projections from orbitofrontal and ventromedial frontal cortex, and the projections from the frontal cortex becoming progressively closer to motor cortex moving dorsolaterally though the striatum (Haber, 2003). This organization has provided insight into the integration of information as required for new learning (Bar-Gad & Bergman, 2001), as well as the development of deficits in progressive diseases such as Parkinson’s Disease, where the expression of motor and cognitive symptoms was considered in accordance to dopamine deficiencies along the anatomical gradient of the striatum (Jokinen et al., 2013; Kish, Shannak, & Hornykiewicz, 1988). With this presumed anatomical basis, the functional gradient mapping of the striatum was examined as a whole in an elegant analysis of resting-state fMRI data (Marquand, Haak, & Beckmann, 2017). Marquand and colleagues (2017) demonstrated the diagonal organization in the human brain and showed that the integrity of this gradient is associated with individual differences in flexible behavior. These findings provide further support for the diagonal organization as the primary topographic principle of the striatum. However, despite the strong positive evidence that suggests a single functional connectivity gradient across the striatum as a whole, the assumption of a unitary functional gradient across the striatum remains to be tested.

We conducted a set of multivariate multiple regressions that can be easily applied to many approaches that attempt to derive or test functional gradient organizations across anatomical space. We apply these models to the striatum to demonstrate their practical utility due to the relatively well-studied striatal anatomy in nonhuman primates, replicating the same results across different human resting-state fMRI samples and across species. The strongest evidence across all test samples was for the model with multiple differentiable gradients within each subdivision of the striatum, the caudate nucleus and putamen, instead of a single gradient across the two nuclei.

## Methods

### Data samples

We utilized data from 4 separate resting-state data samples: the Cambridge Buckner subset of the 1000 functional connectomes, NKI/Rockland data sample, Rockland Enhanced data sample, and Croxson Phillips subset of PRIME-DE. The first three are resting-state fMRI data from human participants, and the fourth is from monkeys. The Cambridge Buckner and NKI/Rockland data samples represent traditional resting-state fMRI protocols while the Rockland Enhanced data sample represents a high resolution resting-state fMRI protocol using multi-band acceleration. Due to various differences in the data and legacy processed data, some of the data samples were processed in slightly different ways as described below.

For the Cambridge Buckner data sample there were a total of 198 human subjects (123 female from the 1000 Connectomes Project (Biswal et al., 2010) (http://fcon_1000.projects.nitrc.org), ages 18-30 (M = 21.03, SD = 2.31). For the NKI/Rockland data sample (http://fcon_1000.projects.nitrc.org/indi/pro/nki.html), there were a total of 207 human subjects (87 female), ages 4-85 (M = 35.00, SD = 20.00). For the Enhanced Rockland data sample, we used the first 377 human subjects (238 female), ages 8-85 (M = 42.11, SD = 20.34). With the Croxson Phillips data sample, there were 9 monkey subjects, but only 6 (5 Macaca mulatta, 1 Macaca fascicularis) with resting state fMRI data (1 female), ages 3.7-8.0 (M = 5.05, SD = 1.56). All subjects in the Cambridge Buckner database passed the screening for motion other artifacts (with at least 2/3 usable data, see below); 18 subjects were excluded from the NKI/Rockland database, leaving 189 subjects (78 female) remaining, ages 4-85 (M = 35.70, SD = 19.89), 78 subjects were excluded from the Rockland Enhanced databased, leaving 299 subjects (194 female), ages 8-85 (M = 40.96, SD = 20.33); and all 6 monkeys with resting state data passed motion artifact screening.

### MRI Acquisition Parameters

Cambridge Buckner data (Siemens 3T Trim Trio): T1-weighted images were collected with MPRAGE with the following image parameters: slices = 192, matrix size = 144 × 192, voxel resolution = 1.20 × 1.00 × 1.33 mm^3^. Resting state fMRI data were T2*-weighted images acquired using EPI with the following parameters: 47 interleaved axial slices, TR = 3000 ms, voxel resolution = 3.0 × 3.0 × 3.0 mm^3^ (119 volumes in total).

NKI/Rockland data (Siemens 3T Trim Trio): T1-weighted images were collected using MPRAGE with the following parameters: slices = 192, matrix size = 256 × 256, resolution = 1.00 × 1.0 × 1.0 mm^3^. Resting state fMRI were acquired with the following parameters: 38 interleaved axial slices, slice gap = 0.33 mm, TR = 2500 ms, TE = 30 ms, Flip Angle = 80 deg, voxel resolution = 3.0 × 3.0 × 3.0 mm^3^ (260 volumes in total).

Enhanced Rockland (Siemens 3T Trim Trio): T1-weighted images were collected using MPRAGE with the following parameters: slices = 176, matrix size = 250 × 250, resolution = 1.00 × 1.0 × 1.0 mm^3^. Resting state fMRI data were acquired with the following parameters: Multi-band Acceleration Factor = 4, 64 interleaved axial slices, slice gap = 0 mm, TR = 1400 ms, TE = 30 ms, Flip Angle = 65 deg, voxel resolution = 2.0 × 2.0 × 2.0 mm^3^ (404 volumes in total).

Croxson Phillips (Philips Achieva 3T): 3 T1-weighted scans were collected with the following parameters: slices = 176, matrix size = 250 × 250, resolution = 0.5 × 0.5 × 0.5 mm^3^. Resting state fMRI data were acquired with the following parameters: 40 interleaved axial slices, TR = 2600 ms, TE = 19 ms, voxel resolution = 1.5 × 1.5 × 1.5 mm^3^ (988 volumes in total).

### Image Preprocessing

Prior to analysis images were preprocessed utilizing SPM12 (http://www.fil.ion.ucl.ac.uk/spm/software/spm12/). For each individual in the Cambridge Buckner and NKI/Rockland data samples, the functional images were first corrected for slice timing, and then realigned to the middle volume according to a 6 parameter rigid body transformation. Structural images were coregistered with the mean functional image, segmented, and then normalized to the MNI template using both linear and nonlinear transformations. Functional images were then normalized utilizing the same parameters as the structural normalization. For Rockland Enhanced, the functional images were also unwarped during realignment to reduce movement related distortions in the data. For the Croxson macaque data, the three T1 volumes were averaged prior to segmentation to increase SNR, and because fieldmaps were available for all macaques, fieldmap correction was applied to the data prior to slice timing correction to reduce distortions, unwarping was performed during realignment, and data were normalized using the National Institute of Mental Health (NMT) macaque template (Seidlitz et al., 2017).

Further preprocessing steps were performed following the standard procedures of resting-state fMRI analysis either using CONN (Whitfield-Gabrieli & Nieto-Castanon, 2012) or custom Matlab scripts. A nuisance regression was constructed with the following confounding variables: 6 motion parameters up to their second derivatives, scans with evidence of excessive motion (Framewise Displacement > .5 or DVARS > 5), effects of session onset, modeled physiological signal generated through aCompCor (Behzadi, Restom, Liau, & Liu, 2007) of the white matter and CSF voxels, and a linear drift component. For the Cambridge Buckner and NKI/Rockland data samples, the residuals of this regression were filtered utilizing a bandpass between the frequencies of 0.008 and 0.09 Hz, while for the other data samples the filtering and nuisance regression were done simultaneously (Hallquist, Hwang, & Luna, 2013). Finally, the resultant data were despiked using a tangent squashing function.

For the macaque data, after the data cleaning procedure, each dataset was split into 5-min time bins, and analyses were run on both the full time-series and each time bin.

### Spatial Gradient Estimation and Model Fitting

An atlas based mask of the right striatum, including both the caudate and putamen as two subdivisions or regions of interest (ROI), was aligned and resampled to the space of each data sample: AAL (Tzourio-Mazoyer et al., 2002) for humans and D99 for macaques (Reveley et al., 2017) (Fig. 1A top, Fig. 1B). We only used the right hemisphere for simplicity, though the results are consistent in the left hemisphere. The cleaned time-series from each voxel within the entire striatum was extracted from each subject and all pairs of correlations were computed, producing a voxel-voxel correlation matrix **R**. While previous methods utilize the similarity matrix generated from a voxelwise fingerprint (Haak et al., 2017), we used the direct time-series correlation between voxels within the striatum for simplicity of computation and interpretation. We find these two methods to approximate each other at high numbers of voxels in the brain (see Fig. S1 for simulations and empirical support).

**Figure 1:**
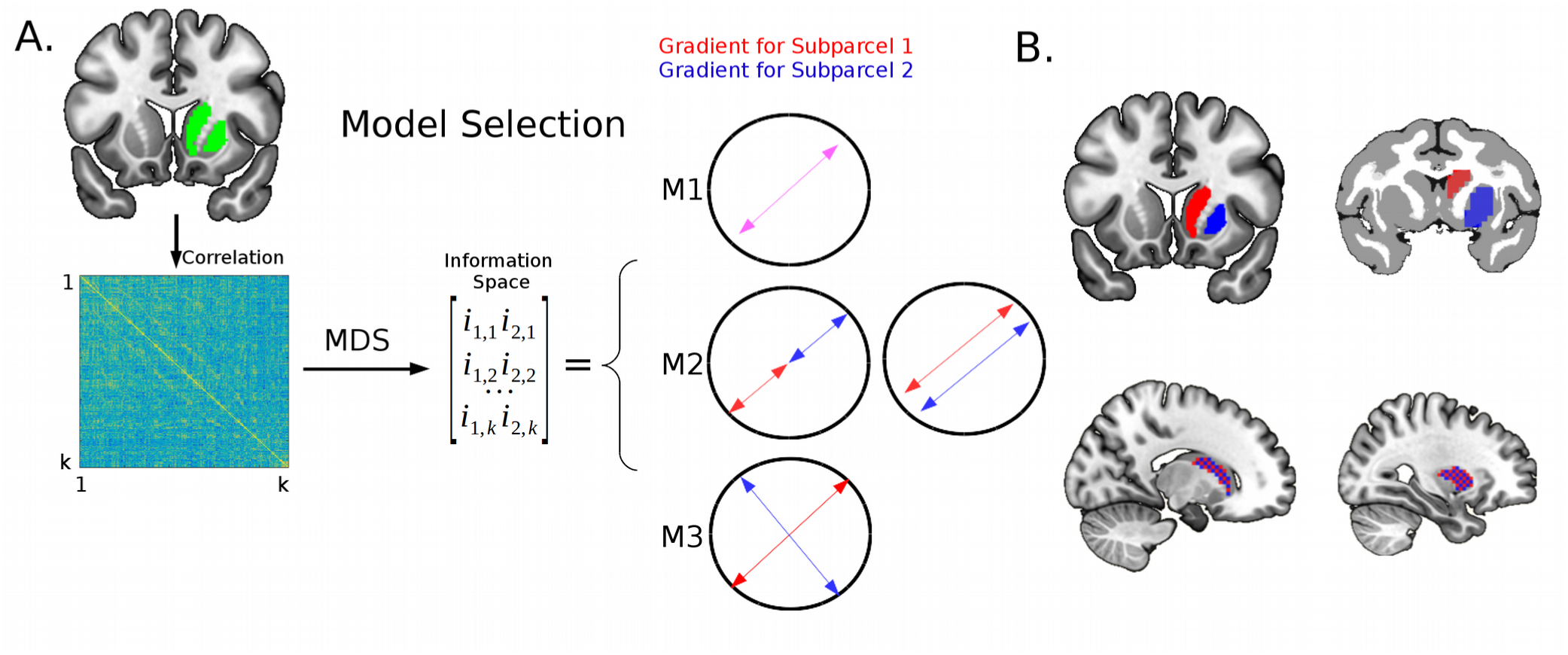
Analysis and Models. **(A)** The time-series of each voxel within the superparcel (caudate + putamen; top left, green) was extracted, and the correlations between all pairs of voxel time-series were calculated. This produced a k by k correlation matrix, where k is the number of voxels within the superparcel. A distance matrix was computed by taking the complement of this correlation matrix, and this distance matrix is reduced into a two dimensional space using nonclassical mutidimensional scaling (MDS), producing an “information space”. To test the three potential gradient models, we then fit this information space with the anatomical space information from the superparcel (i.e. Model 1 [M1]), with Model 2 (M2) including a categorical term (ROI, being within the caudate or the putamen subdivision for each voxel), and Model 3 (M3) also including the interaction between the spatial information and the categorical variable. The bidirectional arrows represent the hypothetical functional gradient(s) within the structure for the case of each winning model. **(B)** Striatal subdivisions for different model testing procedures. **(TOP)** For the striatal model testing procedure in humans, the AAL masks for the caudate (red) and putamen (blue) were eroded, the superparcel being defined as the combination of both. In macaques, the D99 atlas masks for caudate (red) and putamen (blue), with the superparcel being defined as the combination of both. **(BOTTOM)** Caudate (left) and Putamen (right) masks for the checkerboard control analyses.

We then calculated a distance matrix as the complement of the correlation matrix, 1-**R**, such that two voxels with perfect correlation are represented with the minimum distance of 0, and voxels perfectly negatively correlated are represented with a maximum distance of 2. Then, using nonmetric multidimensional scaling (MDS), we constructed a 2-dimensional space that best preserves each voxel’s bivariate rank distance, defined as the space that minimizes the stress of the configuration (the root mean squared difference between the rank order of the distance matrix and the estimated rank order of the distances within the constructed dimensions) (Kruskal, 1964), the result of which we refer to as “information space”. We chose nonmetric MDS because its small set of free parameters (just the dimensionality of the resultant space, which for consistency with prior literature we set to 2 allowing for the estimation of 2 overlapping gradients across space) and allowance for monotonic nonlinearity in the information space (in comparison to the strict linearity metric MDS). However, for demonstration of consistency with the literature, we also used Laplacian Eigenmaps, which constructs dimensions that reproduce the local distances between connected elements of a graph, similar to spectral clustering (Belkin & Niyogi, 2002), with the parameters: {nearest neighbors parameter k = 12 (determines connectivity), heat kernel parameter t = 1 (determines weight of connectivity)}. We found very similar results to those presented in the main manuscript (Fig. S2).

### Model Fitting and Model Selection

To select and fit the appropriate model of the functional connectivity gradients, smooth changes in functional connectivity across the space of the striatum, we applied 3 different multivariate multiple regression models, fitting the anatomical space to the information space (Fig. 1A). Model 1 (M1) predictors contain just the anatomical space information ([x,y,z], rotated with PCA to preserve the spatial variance and mean centered for each striatal subdivision, such that a shift in gradient isn’t encoded by the spatial coordinates themselves):

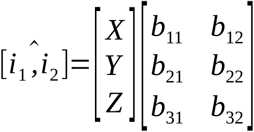

Model 2 (M2) had an additional ROI categorical variable, allowing for a mean shift the spatial gradient across information space for each subdivision:

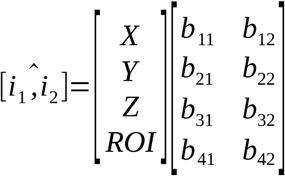

Model 3 (M3) additionally had each interaction between the anatomical space information and the ROI categorical variable, allowing for the gradient the change in slope as well across the subdivisions:

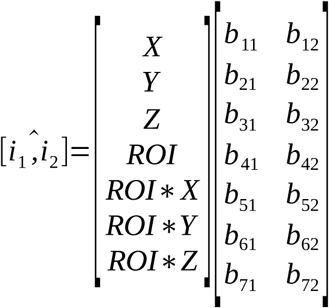

See Fig. 2 for a representation of this analysis in a typical subject, with the anatomical space visualized across information space. The sum of the apparent fits across space in the x, y, and z panes represents the spatial gradient, with the different models allowing the fits for each ROI to either shift or change slope.

**Figure 2:**
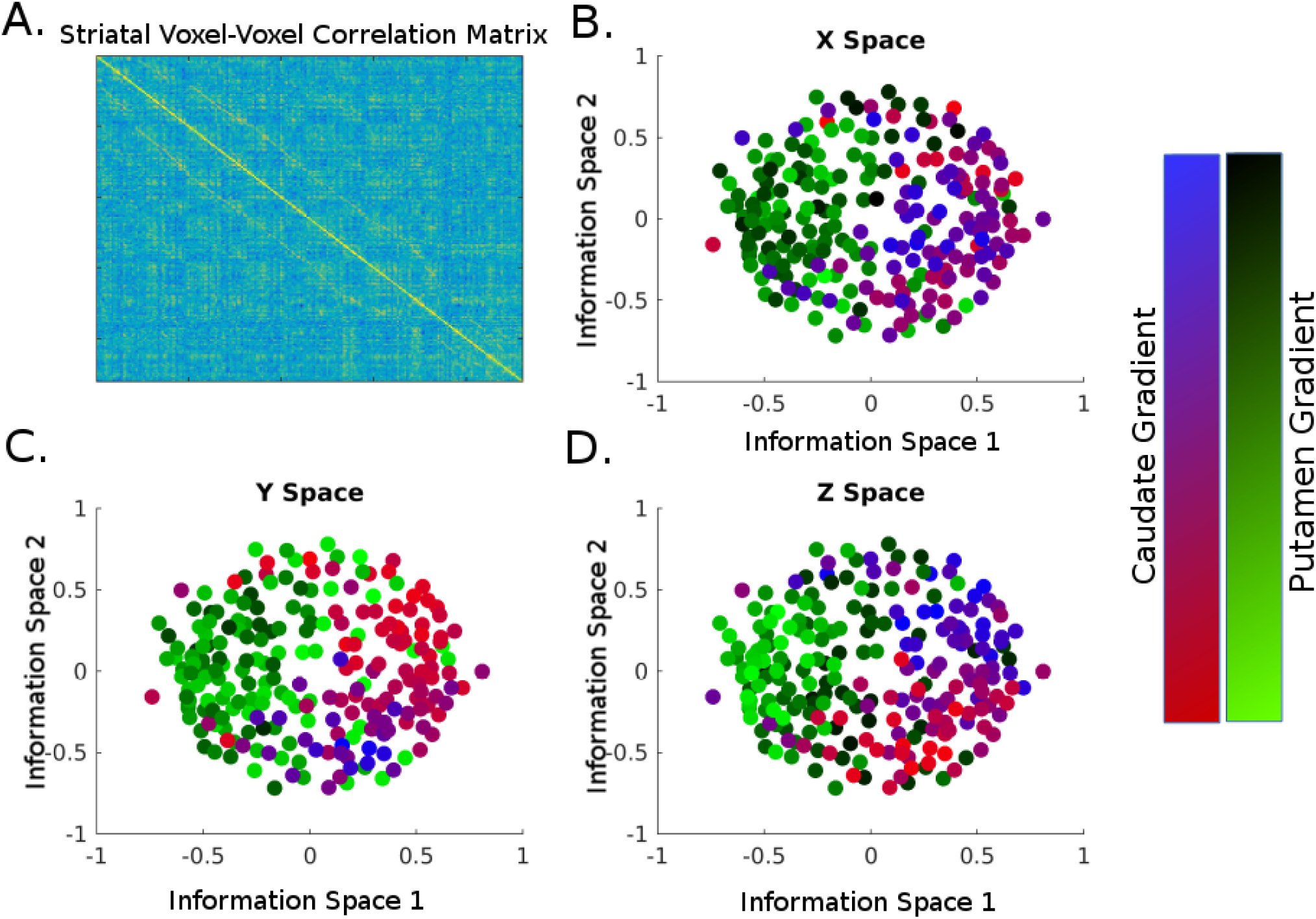
An example subject from the NKI/Rockland database. **(A)** The voxel-voxel correlation matrix from which the information dimensions were subsequently derived. The 3 panes show all voxels in their information space, with the anatomical space (X, Y, Z) of each subdivision or ROI (caudate and putamen) represented as a color gradient, in **(B)**, **(C)**, and **(D)** respectively. A smooth transition of colored dots represents good fit of that anatomical gradient to the information space. Models 1-3 predict different relationships between the coloring of the dots and their position in information space. Model 1, for instance, predicts the red-blue dots to be completely intermingled with the green-black dots, while Models 2 and 3 predict a clear separation between them.

To arbitrate the best fitting model, we used the Bayesian Information Criterion (BIC), kln(n) – 2ln(L), where n is the number of voxels, k is the number of parameters, and L is the maximized likelihood of the model. The BIC provides a rough estimation of the log Bayes Factor, and thus is a convenient substitute for model evidence when priors are unknown or hard to quantify (Kass & Raftery, 1995). This model fitting procedure was done for each subject individually, and the reported values are the average difference in BIC across models (ΔBIC) and proportion of the sample which minimize BIC for each model.

The model selection procedure allows us to select between among the following three possible connectivity gradient patterns within the data: Model 1, the striatum contains a single unitary gradient overlapping across caudate nucleus and putamen, Model 2, the caudate nucleus and putamen contain either a continuation of a single gradient or two parallel gradients (carrying different information in the same direction in space), or Model 3, the caudate nucleus and putamen contain completely different gradients (see Fig. 1B for striatal parcels).

Further, we conducted a series of control analyses to ensure that the model selection procedure was arbitrating the proper models. To do so, we took each ROI and generated checkerboard ROIs for each, generating two subparcels that should produce a single completely overlapping gradient and therefore support Model 1 (Fig. 1B).

If Model 2 is the winning model, the result is ambiguous, as a good fit can come from two possibilities: the mean shift due to ROI can follow along the overall slope of the gradient, or the mean shift can occur orthogonally (Fig. 1A). In the first case, the gradient flows continuously from one ROI into the other. The second case, however, denotes that the information carried along the gradient may be different, in which case we should be reluctant to collapse across the subparcels. To discriminate between these two possibilities, we have developed a statistical procedure to detect nonorthogonal shifts. For each spatial dimension (x, y, and z) we bootstrap the dot product of the two dimensional vector representing the overall gradient and the two dimensional vector representing the ROI shift, e.g.:

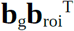

where **b**_g_ = [b_11_,b_12_], **b**_roi_ = [b_41_,b_42_] for the x spatial dimension in Model 2

If the 95% confidence interval of this bootstrapped distribution does not include 0, we can conclude that we have evidence for non-orthogonality in the shift, suggesting a shift along the slope of the overall gradient (allowing for the collapsed analysis) (see Fig. S3 for example subjects).

### Visualization of Gradient Results

To visualize the two estimated spatial gradients (labeled dimension 1 and dimension 2) across the whole striatum, the anatomical space was projected across the gradients, **Xb** mathematically, and plotted onto each voxel within the striatum. Because each subject could theoretically have a different gradient, and even if they follow the same gradient the sign could be arbitrarily flipped, we clustered the gradients prior to averaging across subjects. We applied k-means, selecting the cluster solution (out of cluster solutions 1-10) that maximized the ratio of between cluster distance and within cluster distance (Calinski-Harabasz criterion) (Caliński & Harabasz, 1974) for each striatal subdivision separately. We then averaged all subjects’ estimated gradient projection within cluster in order to avoid heterogeneous averaging.

## Results

### Model Testing of Human Resting State fMRI

Table 1 shows the model fit results for the three models under consideration across different resting-state fMRI data samples, with 95% confidence intervals around ΔBIC providing a classical tests of group based model selection (Stephan, Penny, Daunizeau, Moran, & Friston, 2009).

**Table 1:** Results of model testing procedures

| Experiment | $\overline{\Delta BIC}$ (M1-M2)<br>[95% CI] | $\overline{\Delta BIC}$ (M1-M3)<br>[95% CI] | $\overline{\Delta BIC}$ (M2-M3)<br>[95% CI] | Prop. Support<br>for M1 | Prop. Support<br>for M2 | Prop. Support<br>for M3 |
| --- | --- | --- | --- | --- | --- | --- |
| *Striatal Tests* |  |  |  |  |  |  |
| Cambridge Buckner (n = 198) | 32.17<br>[27.46,36.87] | 23.52<br>[18.09,28.96] | -8.64<br>[-6.50,-10.79] | 0.14 | 0.60 | 0.26 |
| NKI/Rockland (n = 189) | 78.02<br>[68.81,87.23] | 78.16<br>[67.59,88.73] | 0.14<br>[-3.15,3.42] | 0.08 | 0.52 | 0.40 |
| Rockland Enhanced (n = 299) | 104.29<br>[89.62,118.95] | 119.76<br>[102.73,136.80] | 15.47<br>[10.76,20.19] | 0.12 | 0.33 | 0.55 |
| Croxson Macaque (n = 6) | 133.06<br>[-47.46,313.57] | 189.18<br>[-18.99,397.35] | 56.12<br>[22.58,88.66] | 0.00 | 0.00 | 1.00 |
| *Caudate Checkerboard* |  |  |  |  |  |  |
| Cambridge Buckner (n = 198) | -8.55<br>[-8.68,-8.42] | -33.51<br>[-33.82,-33.21] | -24.97<br>[-25.23,-24.70] | 0.99 | 0.01 | 0.00 |
| NKI/Rockland (n = 189) | -8.52<br>[-8.64,-8.40] | -32.86<br>[-33.20,-32.52] | -24.34<br>[-24.65,-24.02] | 1.00 | 0.00 | 0.00 |
| *Putamen Checkerboard* |  |  |  |  |  |  |
| Cambridge Buckner (n = 198) | -8.88<br>[-8.99,-8.77] | -34.87<br>[-35.13,-34.60] | -25.99<br>[-26.23,-25.75] | 1.00 | 0.00 | 0.00 |
| NKI/Rockland (n = 189) | -8.97<br>[-9.07,-8.88] | -34.76<br>[-35.06,-34.46] | -25.78<br>[-26.06,-25.51] | 1.00 | 0.00 | 0.00 |

In the Cambridge Buckner dataset, there was the highest support for Model 2. Further, bootstrapping of the dot product of the vector representing the overall gradient and the vector representing the shift of that gradient across the striatal subdivisions for each subject revealed that for 91.53% of participants that demonstrated preference for Model 2, the shift wasn’t detectably non-orthogonal in at least one of the spatial dimensions (Note: 34.75% having no detectable non-orthogonal shifts in any of the 3 dimensions) (see Fig. S3). For the NKI/Rockland dataset, we found equivalent support for Models 2 and 3. In the higher spatial resolution Enhanced Rockland data sample, we found the highest support for Model 3, indicating differential gradients across the striatal subdivisions.

For validation of the model selection process, model testing with the two checkerboard subparcels of the caudate produce support for a completely overlapping gradient (Model 1), as expected. Similarly, model testing with the two checkerboard subparcels of the putamen also produced the most support for the expected completely overlapping gradient (Model 1).

### Model Testing of Macaque Resting State fMRI

We conducted the same model testing procedures in resting-state fMRI data from 6 macaque monkeys. Model 3, using the same procedures as the prior analyses, demonstrated the highest evidence, replicating the Rockland Enhanced results in these macaque data. We further confirmed the model fitting results separately in each monkey across the 10 time-bins and the proportion winning model by time-bin was calculated for each monkey; the means and standard deviations are 0.10 (0.17), 0.15 (0.19), 0.78 (0.17), for Model 1, Model 2, and Model 3 respectively.

### Clustering and Characterization of the Functional Gradients

To visualize the model fitting of the functional gradients within the striatum and its two subdivisions, we projected the space of the caudate and putamen onto their estimated gradients for each subject within the Enhanced Rockland data sample and plotted it onto the striatum. The Enhanced Rockland data sample were used here for clustering and visualization because of the large number of voxels would make the model estimation more reliable, in addition to the cross species convergence described above. We clustered the spatial gradients of the caudate and putamen separately as Model 3 was the best fitting model. The clustering analysis showed a clear peak in this criterion at k = 4 for the caudate subdivision (Fig. 3). The subjects within a cluster were averaged to produce the 4 gradient maps, one per cluster (Fig. 4). The two predominate gradient organizations were a medial-lateral and a diagonal (from dorsolateral to ventromedial), with participants more commonly having both organizations rather than having one single organization across both estimated spatial gradients (dimension 1 and dimension 2) (Fig. 4B, sum of the main diagonal (↘) vs sum of the anti-diagonal (↗) of the contingency table). The insets shown in Figure 4A display the diagonal gradient in the same fashion as in previous papers (Marquand et al., 2017), which illustrates the similarity of the current results with their results.

**Figure 3:**
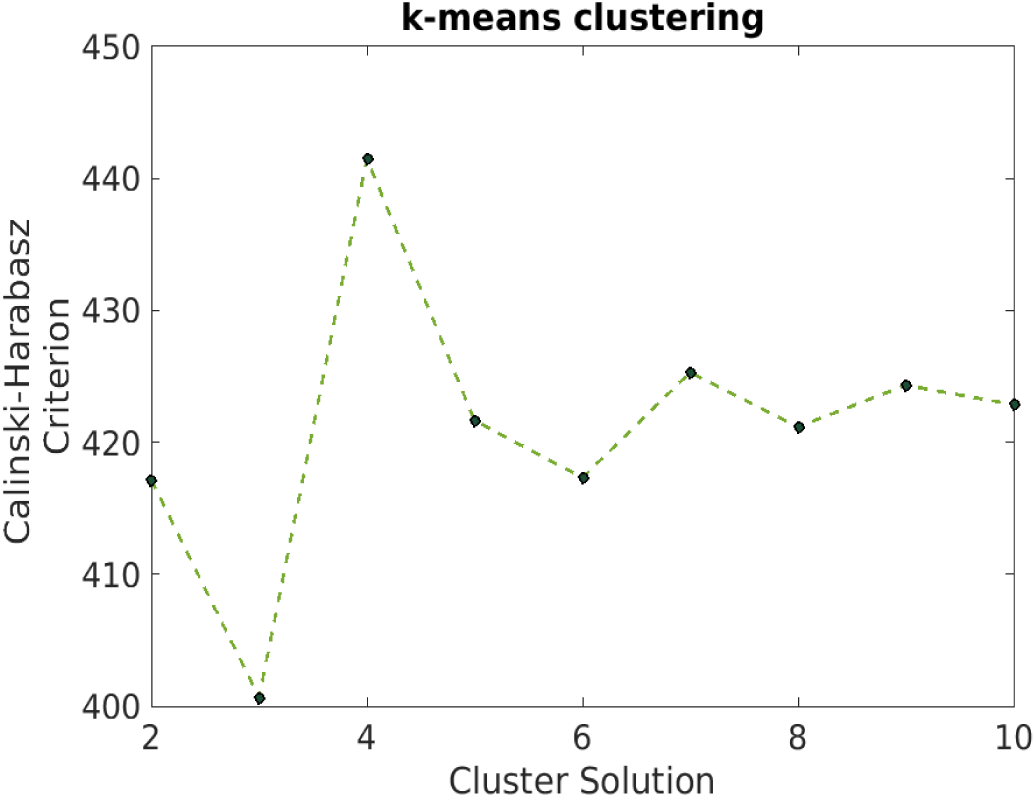
Clustering Solution for the functional gradients of the caudate subdivision. Ratio of between cluster distance and within cluster distance (Calinski-Harabasz criterion) as a function of number of clusters. A clear peak at 4 indicates this as the preferred solution.

**Figure 4:**
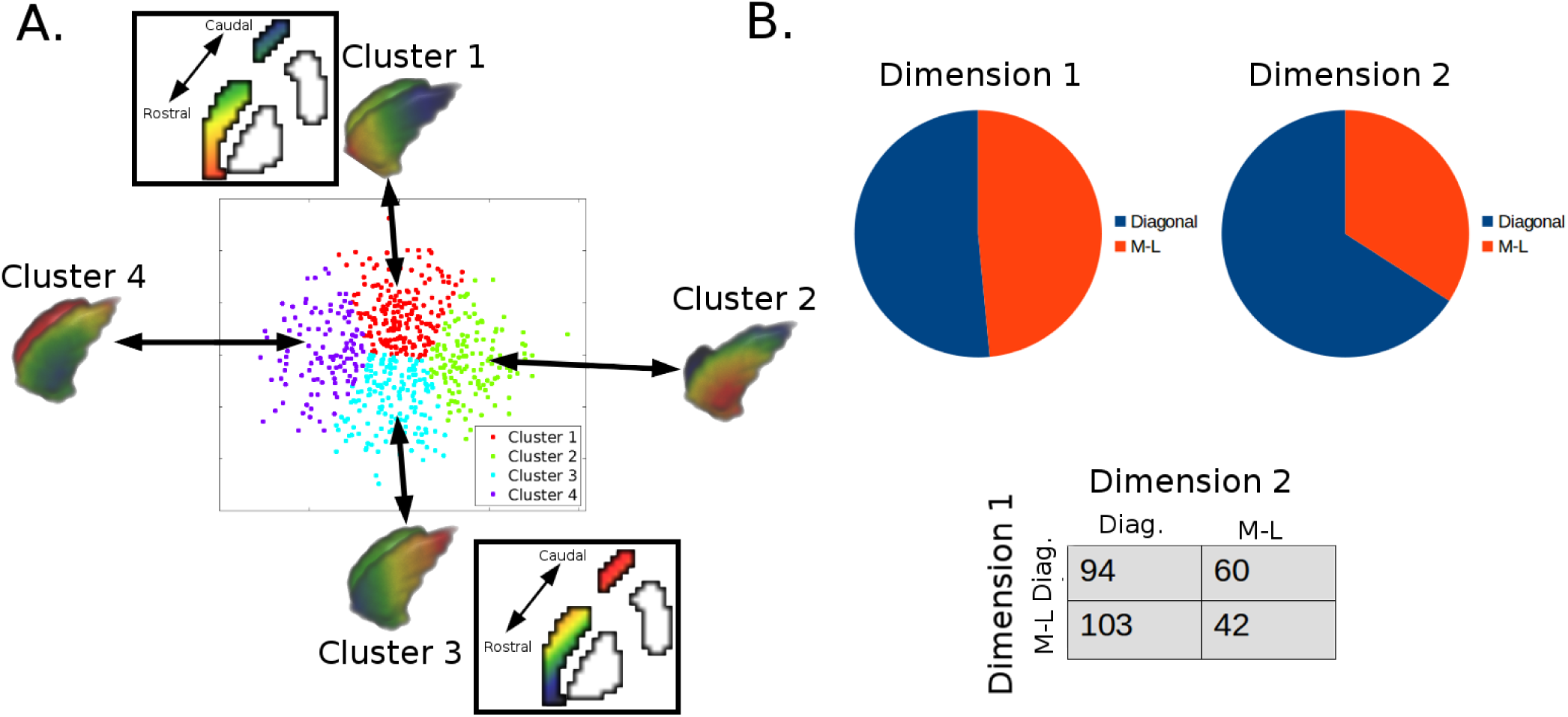
Clustering of the gradient results for the caudate. **(A)** K-means clustering was applied to every subject’s gradients estimated for dimension 1 and dimension 2, visualized by the scatter plot where each dot represents one of these estimated gradients. The mean gradient of each cluster is visualized, clusters 1 and 3 representing the flipped versions of the same diagonal gradient, while 2 and 4 represent flipped versions of the same medial-lateral gradient. The insets next to clusters 1 and 3 are representations of each of these gradients visualized in the same formate as the prior literature. This is to show the similarity in the diagonal gradient between our results and Marquand et al.’s (2017) results. **(B)** Top: proportion of subjects displaying each organization within each estimated gradient. Bottom: A contingency table illustrates that displaying both gradients (anti-diagonal) is more common than displaying a single organization across both estimated gradients (main diagonal)

The same k-means clustering procedure was applied to the spatial gradients of the putamen. The preferred clustering solution was k = 2 (Fig. 5A). This clustering result revealed a diagonal or dorsolateral-ventromedial gradient as the primary organization (Fig. 5B). This result was the same even when forced into a cluster solution of 4.

**Figure 5:**
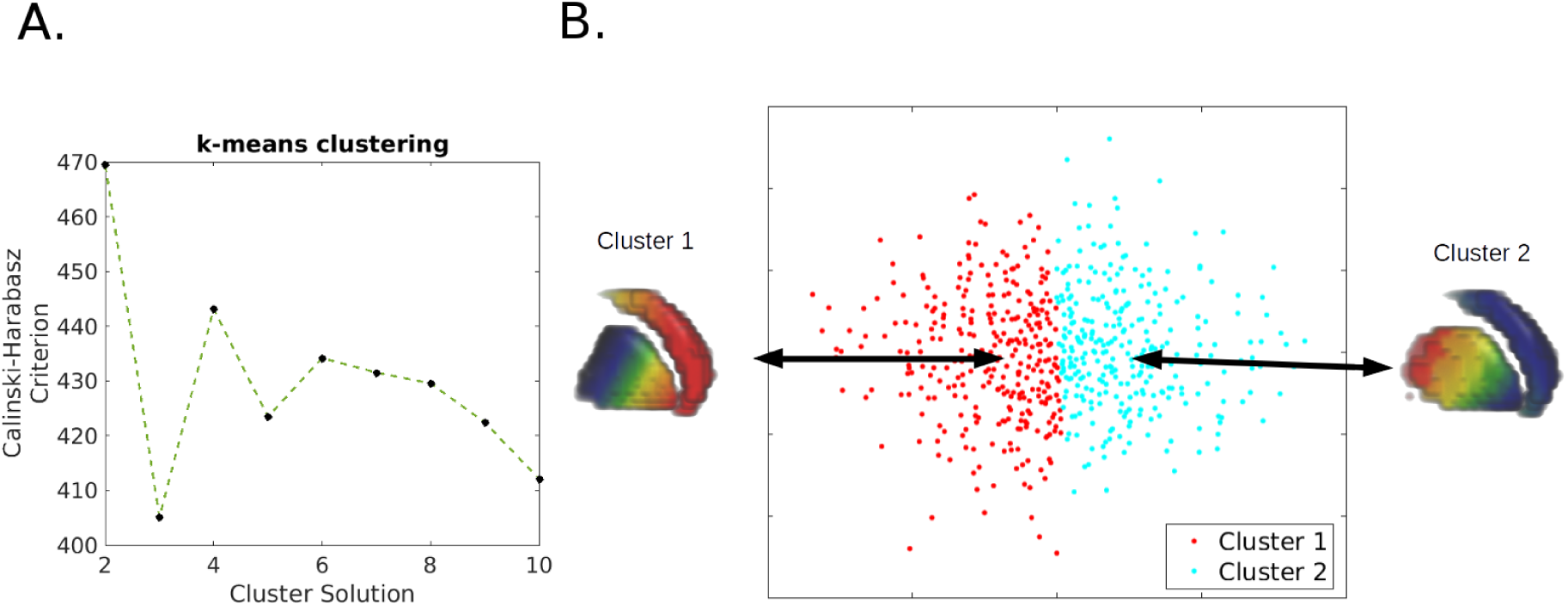
Clustering of the gradient results of the putamen. **(A)** A peak in the ratio of between cluster distance and within cluster distance (Calinski-Harabasz criterion) at k = 2 indicates this as the preferred cluster solution. **(B)** The two cluster solution demonstrates the diagonal gradient as the primary functional gradient for the putamen.

Further, we estimated the spatial gradients while assuming Model 1 with a stable single gradient across both the caudate and putamen subdivisions of the striatum. With clustering based on the caudate gradient, we now found the preferred cluster solution to be k = 2, and forcing k = 4 led to the same conclusion: only the diagonal organization was detected (Fig. S4). This suggests that assuming that a region of interest contains a single spatial gradient organization can obscure other important organizational features if the region actually contains two subregions with different spatial gradient organizations.

Finally, we performed this projection procedure in the macaque sample, demonstrating similarities to the human data, with the caudate demonstrating similarly both diagonal like organizations and medial lateral organizations (Fig. 6). In addition, there seems to be significant individual differences in functional gradient organizations, as depicted in the figure by plotting each individual monkey’s functional connectivity gradients (see Fig. S5 for quantifications).

**Figure 6:**
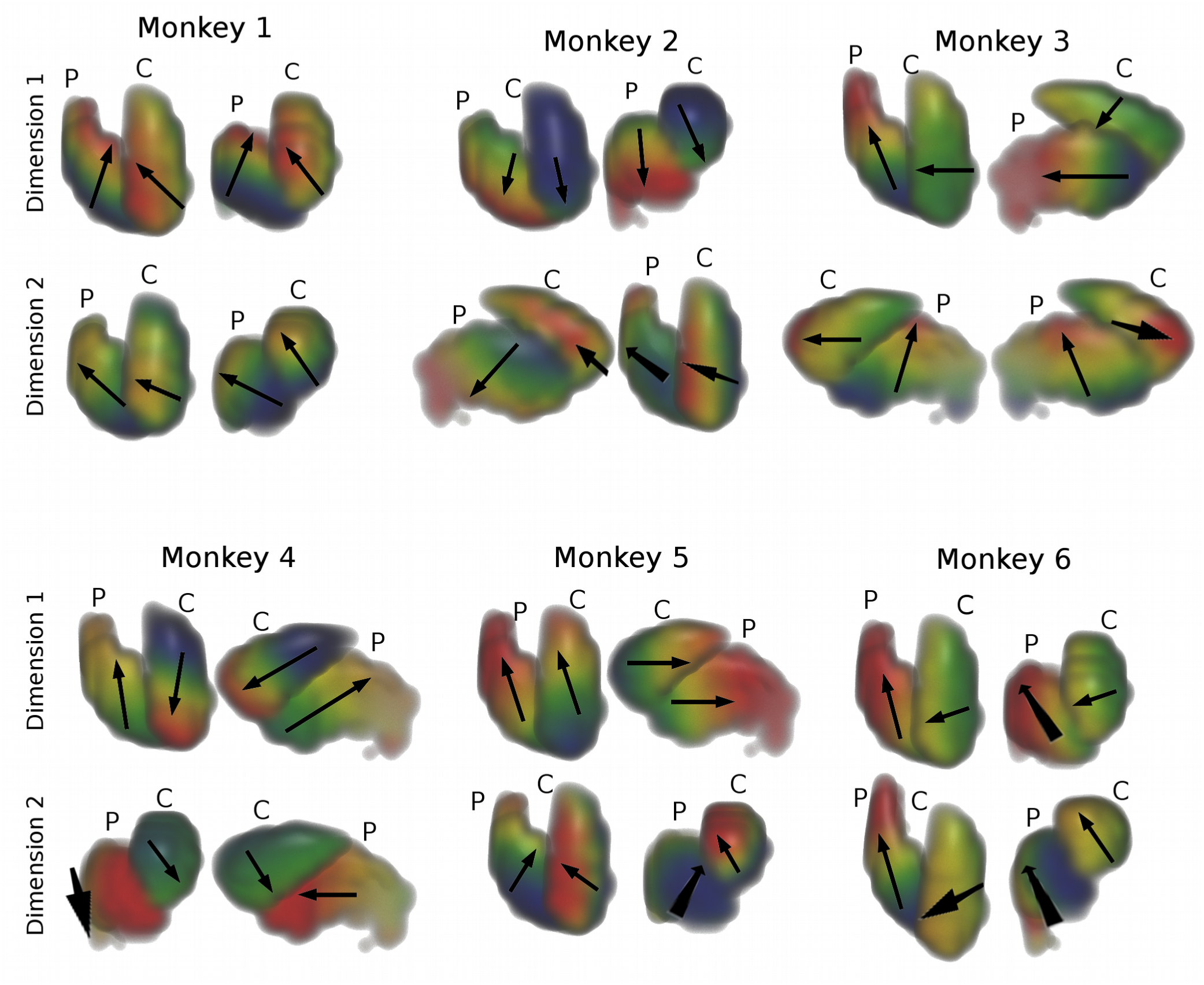
Gradient projection in each individual monkey of the Croxson data sample. The nonhuman primates also showed individual differences in functional topology of the striatum, with monkeys demonstrating one or two motifs out of several. With the caudate, these motifs include medial-lateral (Monkey 2 Dim 2, Monkey 3 Dim 1, Monkey 6 Dim 1), diagonal (Monkey 2 Dim 1, Monkey 3 Dim 2, Monkey 4 Dim 1, Monkey 5 Dim 1), and “strict” diagonal (dorsolateral – ventromedial with no anterior-posterior component) (Monkey 1 Dim 1 & 2,, Monkey 4 Dim 2, Monkey 5 Dim 2, Monkey 6 Dim 2). Consistent with the modeling results, no monkey demonstrated identical gradients within caudate and putamen across both estimated gradients/information dimensions.

## Discussion

An assumption of previous functional connectivity gradient analyses is that there is a unitary gradient (or several overlapping unitary gradients) across the entire space of an anatomical structure. In this paper, we presented a model testing procedure for this assumption that should be performed prior functional connectivity gradient mapping within a parcel. Applying three models to test the gradient organization of the striatum, we demonstrated that even for the case of the striatum, where a single or dominant gradient organization was well supported for the structure, this assumption seems untenable and perhaps overly simplified.

Consistently, across all datasets, Model 1 demonstrated a poor fit to the data in comparison to either Model 2 or Model 3. However, support for Model 2 or Model 3 seemed to vary systematically across databases, with more highly sampled data, across time or space, leading to more evidence towards Model 3. It might be the case that more highly sampled data leads to more accurate estimation, both at the level of the correlation matrix (affected most strongly by time), and at the level of the gradient estimation (affected most strongly by space). This possibility is hard to dissociate from another possibility that might have neural implications: different multiplexed organizations may be observable at different scales. A limitation of this study is a lack of directly comparable data at different resolutions, which may provide an indication as to whether this is the case.

In sum, our model testing procedure showed that the caudate and the putamen, two parcels of the striatum, demonstrate differential gradient organizations instead of a single unitary gradient. This result was replicated across across species. While the primary purpose of this paper was to introduce our approach of systematic testing of the assumptions in spatial gradient analysis, this result requires some consideration. Firstly, the primary support for the single diagonal gradient across both caudate and putamen comes from rigorous anatomical tracing studies in nonhuman primates (e.g. Haber, 2003). While we demonstrated support for Model 3 across human and nonhuman primates, it’s possible that the gradient differences occur solely due to differences in functional connectivity organization obtained from resting-state fMRI data, which may not represent the structural organization shown in the anatomical tracing work. Indeed, direct comparison of human anatomical connectivity through diffusion imaging and human functional connectivity shows relatively weak convergence (Skudlarski et al., 2008) and this convergence can by increased by modeling functional dynamics (Honey et al., 2009). Secondly, it is also possible that the more exhaustive coverage of MRI uncovered a more subtle difference in organization (e.g. Choi et al., 2012) than the sparse sampling approach in tracing studies. Lastly, the generalized diagonal scheme as a unitary gradient is possibly a simplification of the overall underlying organization. For a unitary gradient to account for the data, two qualities of the data must be satisfied: the projections must be similar in nature (origin), and they must terminate in the same relative location along each nucleus. Some of the direct evidence cited for the overall diagonal organization contradicts the plausibility of a single unitary gradient: the sharp reduction in projections from caudal motor cortical areas to the caudate in comparison to the putamen (McFarland & Haber, 2000) and the restriction of extent of DLPFC projections to mostly the rostral putamen while strongly projecting across the entire rostral-caudal axis in the caudate (Selemon & Goldman-Rakic, 1985).

The convergent result of different gradients across the caudate and putamen in this report provides new motivation for investigating the functional organization of previously assumed homogeneous structure. Intriguingly, the second organizational gradient we found within the caudate was a medial-lateral organization. This organization corroborates prior nonhuman primate tracing studies (Selemon & Goldman-Rakic, 1985). Given the delineation of the diagonal organization within humans, and the demonstration of its behavioral relevance (Marquand et al., 2017), it seems that an important next step is to investigate the relevance of this medial-lateral organization in humans.

It’s worth it to note that our general approach to fitting gradients is a bit different than previous work (Haak et al., 2017). We estimate the change in information across space anatomical space linearly. A intuitive description of the spatial fitting step is an attempt to describe the mapping from the anatomical space to information space. While Haak and colleagues (2017) employ a nonlinear mapping approach (similar to a nonlinear warping procedure), the procedure reported here is a simple linear mapping of information space to anatomical space (similar to a rigid body transformation). The approach we took constrains the possible solutions, and thus it should be noted that nonlinear organizations across space will not be detected by our metric. However, the majority of topologies found in neuroscience are described in terms of linear transformations of the space, leaving the approach appropriate for the current application. Indeed, the discovery of the same gradient of organization within the caudate as Marquand et al.’s (2017) (Fig. 4A) further lends credence to the usefulness of this approach.

The striatum as a test case has provided a clear demonstration of the importance of considering the possibility of multiple functional connectivity gradients and the development of model testing procedures to arbitrate between the different possibilities. In sum, the procedure described in this report allows investigators to test three models of functional connectivity gradients within a region of interest. In the simplest case is full support of Model 1, as the parcel should be able to be treated as a single entity. Support for Model 2 is the same in the case where the mean shift in gradients is along the slope of the gradient. However, in cases of support for Model 3, or the case of support for Model 2 where the mean shift is not along the slope of the gradient, the subparcels must be considered separately in the estimation and examination of their gradient mapping.

